## Supplemental Figures for "Loss of ZNRF3/RNF43 Unleashes EGFR in Cancer"

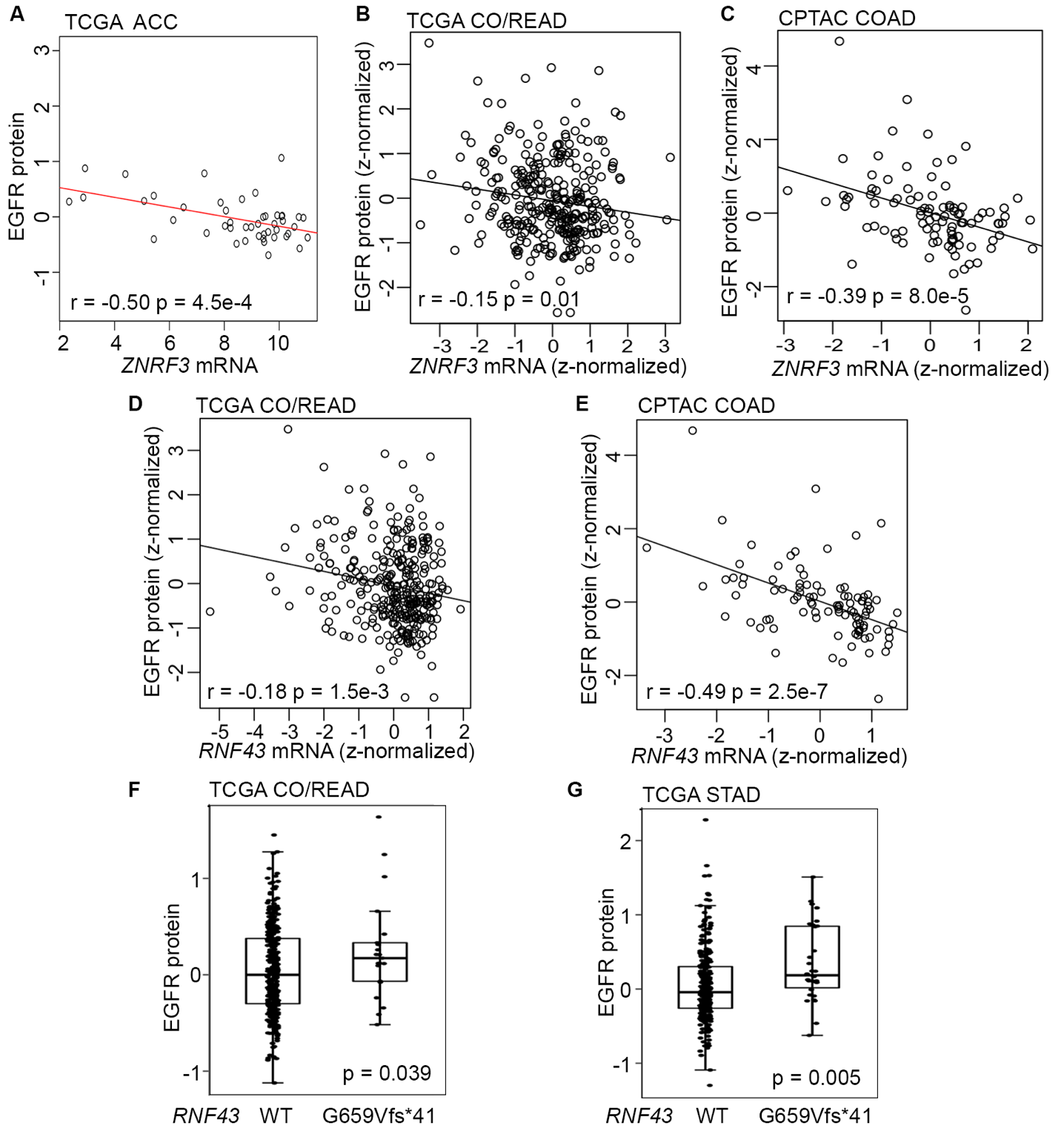

Fig. S1 EGFR protein level is negatively associated with *ZNRF3*/*RNF43* mRNA expression in cancers.

**A**. Scatterplot of EGFR protein level (signal) versus *ZNRF3* mRNA expression (RSEM, Log2 (Val+1)) using the TCGA adrenal cortical carcinoma (ACC) dataset.

**B**. Scatterplot of EGFR protein level (z-normalized) versus *ZNRF3* mRNA expression (z-normalized) using the TCGA colorectal adenocarcinoma (CO/READ) dataset.

**C**. Scatterplot of EGFR protein level (z-normalized) versus *ZNRF3* mRNA expression (z-normalized) using the CPTAC colon adenocarcinoma (COAD) dataset.

**D**. Scatterplot of EGFR protein level (z-normalized) versus *RNF43* mRNA expression (z-normalized) using the TCGA colorectal adenocarcinoma (CO/READ) dataset.

**E**. Scatterplot of EGFR protein level (z-normalized) versus *RNF43* mRNA expression (z-normalized) using the CPTAC colon adenocarcinoma (COAD) dataset.

**F**. Boxplot of EGFR protein levels (signal) in human colorectal adenocarcinomas expressing RNF43 WT or RNF43 G659Vfs*41, using the TCGA dataset.

**G**. Boxplot of EGFR protein levels (signal) in human stomach adenocarcinomas (STAD) expressing RNF43 WT or RNF43 G659Vfs*41, using the TCGA dataset.

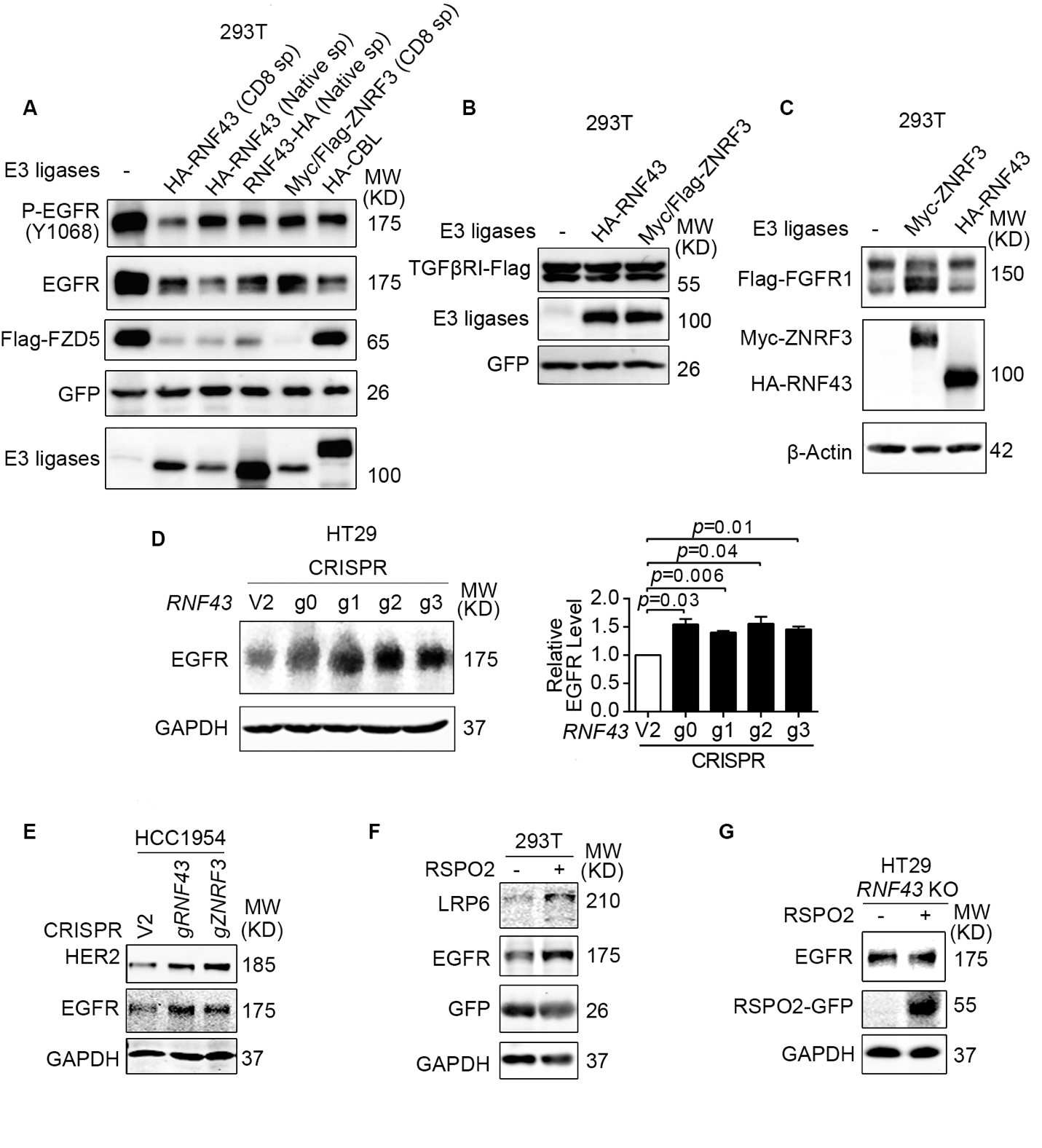

Fig. S2 ZNRF3/RNF43 negatively regulates EGFR protein level.

**A**. Overexpression of RNF43 or ZNRF3 decreases P-EGFR, total EGFR, and FZD5 protein levels in 293T cells. GFP, EGFR, Flag-FZD5 constructs were co-transfected with E3 ligase constructs or vector control. CBL overexpression serves as a positive control for EGFR downregulation. RNF43 or ZNRF3 were expressed with their native signal peptides or CD8 signal peptide.

**B**. Overexpression of RNF43 or ZNRF3 has no impact on TGF-β receptor I (TGFβRI) protein level in 293T cells. TGFβRI-Flag construct was co-transfected with RNF43, ZNRF3 or vector control.

**C**. Overexpression of ZNRF3 or RNF43 does not decrease FGFR1 protein level in 293T cells. Flag-FGFR1 construct was co-transfected with ZNRF3, RNF43, or vector control.

**D**. Expression of Cas9-CRISPR *RNF43* guide RNA enhances EGFR protein level in HT29 cells, shown by representative Western blot images (left panel) and quantification results (right panel). Means ± SEMs are shown. *p*-values were calculated by one-way ANOVA uncorrected Fisher’s LSD test.

**E**. Expression of Cas9-CRISPR *RNF43* guide RNA or *ZNRF3* guide RNA enhances both EGFR and HER2 protein levels in HCC1954 cells.

**F**. RSPO2 treatment (50 ng/ml, 2-4 hr) enhances EGFR and LRP6 protein levels in 293T cells infected with lentivirus expressing FUCGW-EGFR.

**G**. Overexpression of RSPO2 WT does not enhance EGFR protein level in HT29 *RNF43* knockout cells.

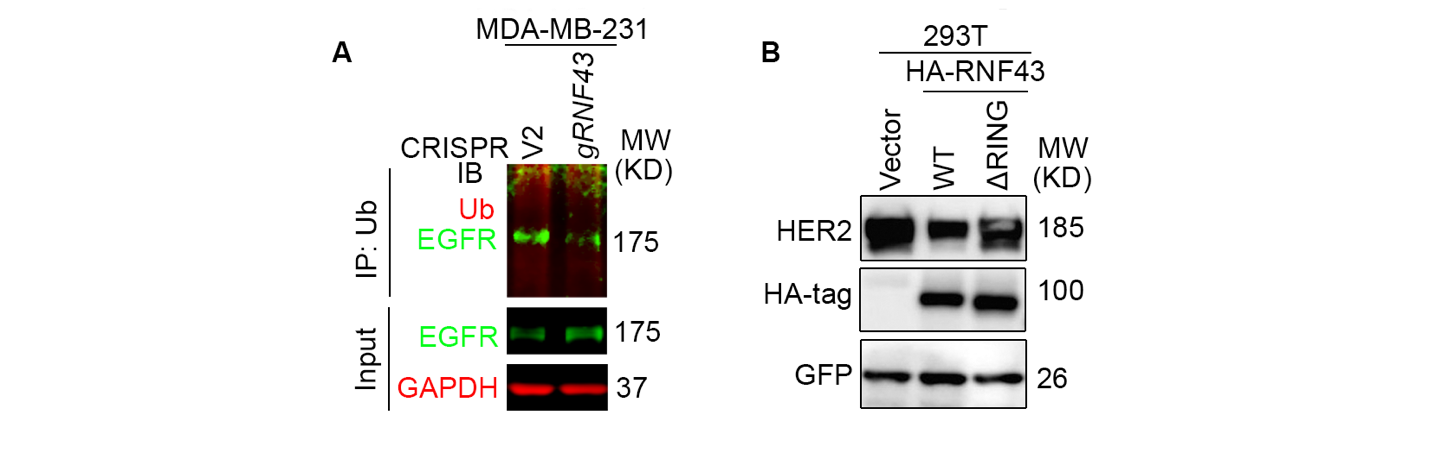
Fig. S3 RNF43 loss decreases EGFR ubiquitination.

**A**. Depletion of *RNF43* by CRISPR/Cas9 decreases EGFR ubiquitination in MDA-MB-231 cells. EGFR ubiquitination was examined by Ub IP followed by EGFR IB.

**B**. RNF43 downregulates HER2 protein level through the RING domain. 293T cells were co-transfected with HER2 and Vector, RNF43 WT or ΔRING mutant.

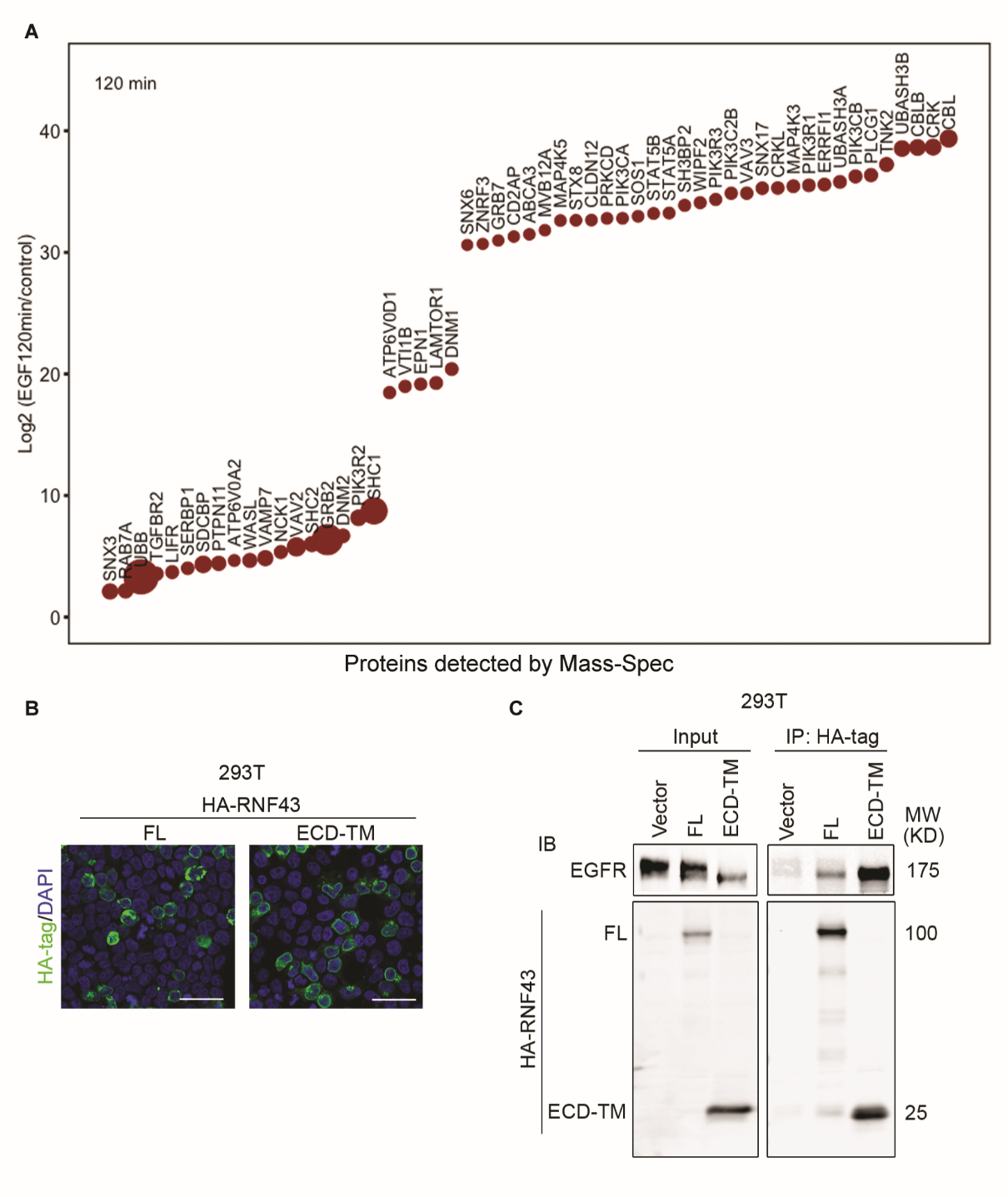

Fig. S4 ZNRF3/RNF43 interacts with EGFR

**A**. Relative abundance of the EGFR-associated proteins. BGC823 cells were starved overnight and then treated with 50 ng/ml EGF for 0 min (control) or 120 mins (EGF-stimulated). Endogenous EGFR was immunoprecipitated by anti-EGFR antibodies. EGFR-associated proteins with over 10^5 iBAQ (intensity Based Absolute Quantification) and over 2-fold increase in iBAQ after EGF stimulation were plotted. The areas of the circles indicate the abundance of iBAQ of EGFR-associated proteins obtained in EGF-stimulated IPs. The y axis indicates the fold change of iBAQ of EGFR-associated proteins identified in EGF-stimulated versus control in the log2 scale, which are arranged from low to high along the x axis.

**B**. Immunofluorescence staining for HA-tagged RNF43 FL and RNF43 ECD-TM in 293T cells transfected with RNF43 constructs. Scale bar=40 μm.

**C**. RNF43 ECD-TM interacts with EGFR. 293T cells were co-transfected with EGFR and HA-tagged RNF43 constructs. EGFR interaction with RNF43 FL and RNF43 ECD-TM were examined by HA-tag IP followed by EGFR IB.

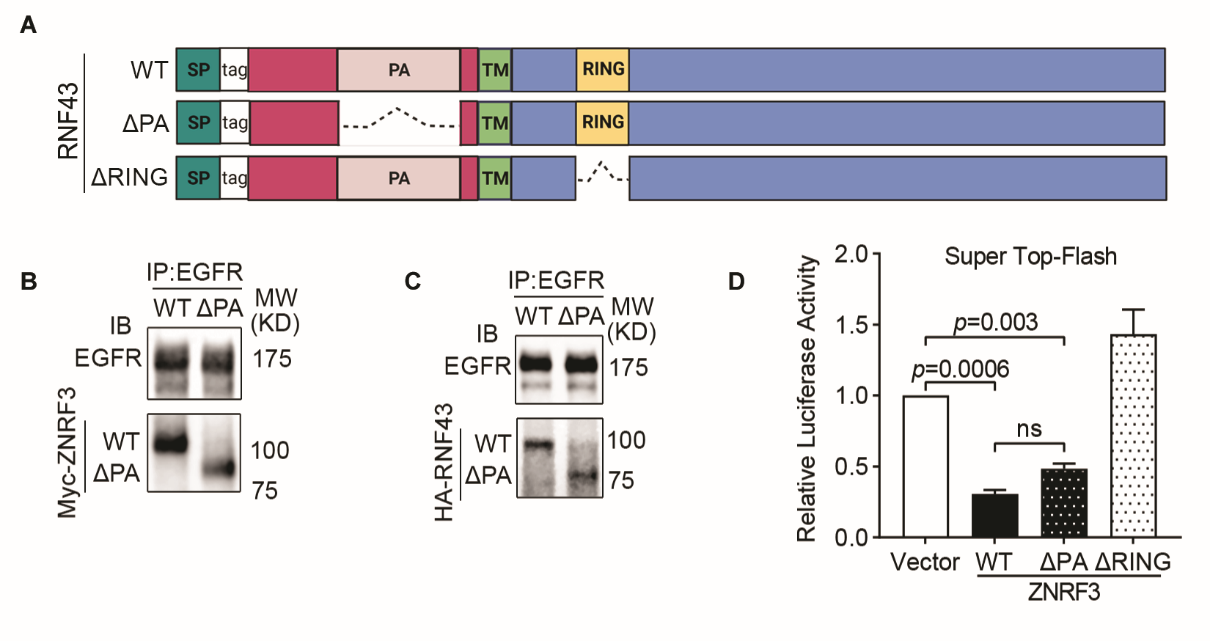

Fig. S5 The protease associate domain of ZNRF3/RNF43 is dispensable for EGFR interaction.

**A**. Schematic diagram of tagged RNF43 wild-type and mutant proteins. SP, signal peptide; WT, wild-type; PA, protease associate domain; TM, transmembrane domain; RING, E3 ligase RING domain.

**B**, **C**. EGFR interacts with the ΔPA mutant of Myc-tagged ZNRF3 (**B**) and HA-tagged RNF43 (**C**). 293T cells were co-transfected with EGFR and ZNRF3/RNF43 WT or ΔPA constructs.

**D**. Overexpression of ZNRF3 WT or ΔPA mutant inhibits canonical WNT signaling in 293T cells in Super Top-Flash assay. ZNRF3 ΔRING mutant serves as a negative control. Means ± SEMs are shown. *p*-values were calculated by one-way ANOVA uncorrected Fisher’s LSD test. n.s., not significant.

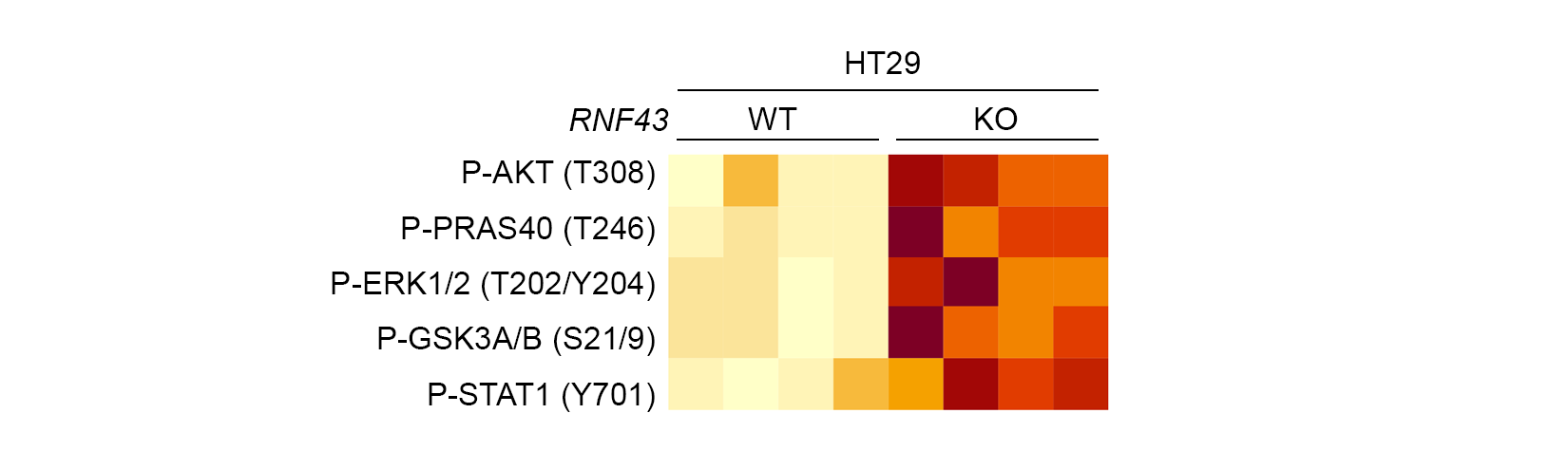

Fig. S6 RPPA identifies EGFR downstream signaling molecules upregulated by *RNF43* knockout in HT29 cells.

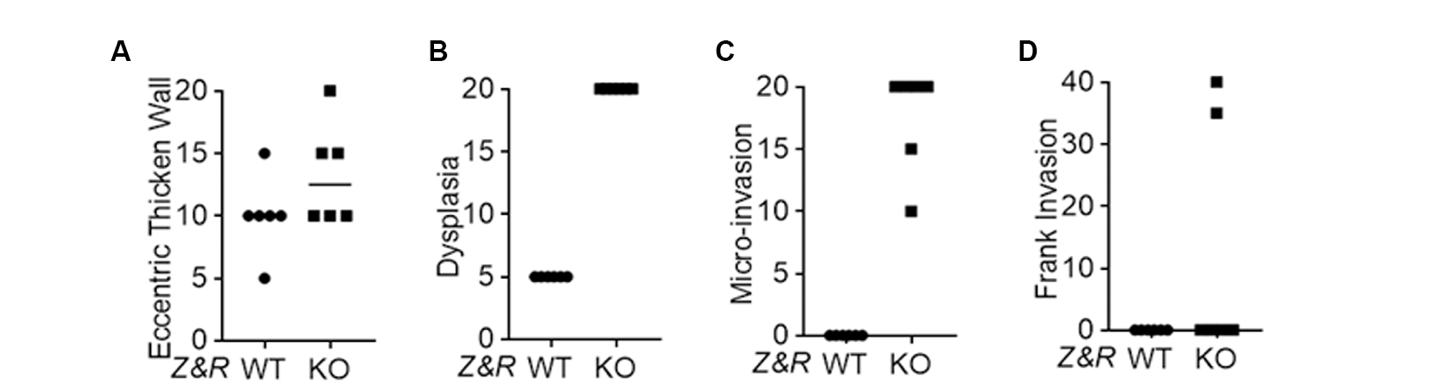

Fig. S7 Pathological assessment on eccentric thicken wall (A), dysplasia (B), micro-invasion (C), and frank invasion (D) of WT and *Znrf3*/*Rnf43* KO mouse prostate tissues.

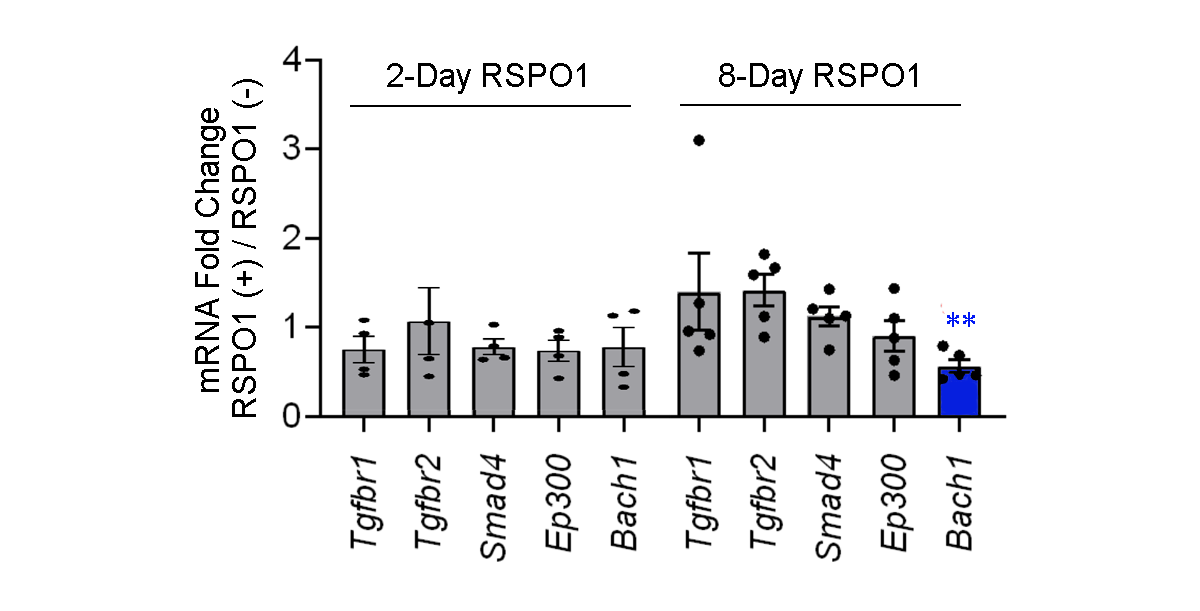

Fig. S8 qPCR analysis for TGF-β signaling relevant genes in *Apc*^min^ mouse intestinal tumor organoids cultured with or without RSPO1 supplements. Genes with no significant changes after RSPO1 treatment were plotted in grey, genes significantly down-regulated after RSPO1 treatment were plotted in blue. Means ± SEMs are shown. Welch’s t-test was used to assess statistical significance. **, *p*-value < 0.01.
